# Assessment of the CTCF Binding Sites and Repeat-Positions Upstream the Human *H19* Gene

**DOI:** 10.1101/250407

**Authors:** Minou Bina

## Abstract

The human *H19* and *IGF2* genes share an Imprinting Control Region (ICR) that regulates gene expression in a parent-of-origin dependent manner. Understanding of the ICR sequence organization is critical to accurate localization of disease-associated abnormalities including Beckwith-Wiedemann and Silver-Russell syndromes. Previous studies established that the ICR of the *H19 - IGF2* imprinted domain included several repeated DNA segments. Using BLAST, BLAT, and Clustal Omega, I conducted detailed sequence comparisons to evaluate the annotation of the unique-repeats upstream of the *H19* transcription start site (TSS) and to investigate the extent of similarities among the various repeats. Initial analyses confirmed the existence of two DNA segments consisting of two types of repeats (A and B). However, I find that one of the repeats (B7) is unlikely to be a partial repeat. I provide the genomic positions of the various repeats in the build hg19 of the human genome. I also evaluated the previously predicted CTCF sites (1 to 7) in the context of the ENCODE data: including the positions of DNase I HS clusters and results of ChIP assays. My evaluations did not support the existence of CTCF site 5. Furthermore, the ENCODE data revealed a previously unknown chromatin boundary (consisting of CTCF, RAD21, and SMC3), in a CpG island (CpG27) between the A1 repeat and the *H19* TSS. Furthermore, a sequence within this boundary corresponds to a newly discovered CTCF site (I named it CTCF site 8). My discovery of this chromatin boundary in CpG27 entails mechanistic implications.

## INTRODUCTION

The Mammalian genomes include a group of genes that are imprinted to impart parent-of-origin-specific gene expression reviewed in (2-4). The human chromosomal band 11p15.5 encompasses an imprinted domain (*H19 - IGF2*) that is regulated by an ICR, at about 2 kb upstream of the *H19* TSS. Nonmethylated ICR drives transcription from *H19* gene in the maternal allele; ICR methylation leads to the expression of the *IGF2* gene from the paternal allele (5). In humans, imprinting disorders cause developmental abnormalities including Beckwith–Wiedemann and Silver–Russell syndromes (6). in Silver–Russell syndrome up to 50 % of methylation defects are in imprinted sequences on chromosome 11p15 (7). Abnormalities associated with Beckwith–Wiedemann syndrome include hypermethylation of the ICR that regulates the *H19 - IGF2* imprinted domain (6, 8).

In investigation of the *H19 - IGF2* ICR, a previous study performed detailed BLAST analyses to define the upstream boundary of the imprinted DNA segment, and to determine whether Wilms tumors with loss of imprinting are biallelically CpG-methylated (1). This study identified several unique repeats. Since the location and annotation of these repeats are extensively used in studies of the human *H19 - IGF2* ICR and developmental abnormalities, I wished to assess the accuracy of the repeats. I also wished to evaluate the ICR sequences implicated in binding the transcription factor CTCF (9, 10). Specifically, it is well known that the ICR of the *H19* - *IGF2* interacts with the transcription factor CTCF to form chromatin boundaries (5, 11), and references therein. Usually, topological domains could arise through interactions among boundaries associated with CTCF and cohesin. Cohesin complexes are produced from SMC3, SMC1, RAD21, STAG1/SA1, and STAG2/SA2 (11-15). In cells, CTCF recruits a subset of cohesin subunits to affect chromatin architecture and gene expression (11,13,15,16). CTCF largely determines the cohesin subunits that associated with transcriptionally active enhancers and promoters (15). Furthermore, cohesin and CTCF shape the organization of topological domains in non-redundant ways: while cohesin is important to shaping the domains, CTCF separates neighboring folding domains and keeps cohesin in place (11).

## RESULTS AND DISCUSSION

### Organization of sequences upstream of the H19 TSS

Previously, upstream sequences of the human *H19* gene were found to contain two reiterated units (1 and 2) consisting of two 450-bp direct repeats (A1 and A2) and several 400-bp repeats (B1 to B7), two of which (B4 and B7) were reported to be incomplete (1). To validate the position and to determine the extent of similarities among the repeats, I obtained their annotations from the GenBank accession number AF125183, reported in reference (1). Initially, I mapped repeat-positions in the build hg19 of the human genome offered at the UCSC genome browser (Table 1). For assessment, I closely examined the relative repeat-positions with respect to genes, the ENCODE data (17), including the position of DNase I hypersensitive sites (HS) and results of chromatin immunoprecipitation assays (ChIPs) reported for CTCF, RAD21, and SMC3 (17, 18).

**Table 1.** genomic positions and length of A and B-repeats

| Repeat type | Genomic position in Build hg19 | ~ Length |
| --- | --- | --- |
| A1 | chr11:2,020,617-2,021,067 | 451 bps |
| B1 | chr11:2021068-2021474 | 407 bps |
| B2 | chr11:2021475-2021880 | 406 bps |
| B3 | chr11:2021881-2022287 | 407 bps |
| B4p | chr11:2022346-2022481 | 136 bps |
| A2 | chr11:2,022,868-2,023,312 | 445 bps |
| B5 | chr11:2023313-2023714 | 402 bps |
| B6 | chr11:2023715-2024120 | 406 bps |
| B7p | chr11:2024163-2024414 | 252 bps |
| B7L | chr11:2024121-2024516 | 396 bps |

B4p refers to B4 partial, B7p to B7 partial, B7L to a nearly complete B7 repeat.

To evaluate the reported repeats (1), I performed extensive sequence similarity searches using BLAST (19), BLAT (20), and Clustal Omega (21). Initially, I analyzed overlapping genomic DNA segments derived from the region between the *H19* and *IGF2* genes. I found that overall results obtained from BLAST were relatively accurate but its pitfalls included outputs that were very labor intensive to analyze. Furthermore, it seemed that BLAST omitted short sections of sequence similarities that could be important. The pitfall of BLAT was that its outputs were bizarre since the program was not designed for analyzing unique repeated DNA. While Clustal O was relatively convenient to use, the program could not handle relatively long sequences. Nonetheless, outputs of Clustal O provided a robust way for closely examining the extent of sequence similarities between two or several aligned sequences.

With Clustal O, initially I assessed the relative repeat positions, by comparing the nucleotide sequences of two DNA sections that I downloaded from the UCSC genome browser: one section encompassed A1, B1, B2, B3, and B4; the other A2, B5, B6, and B7 (Appendix A). For evaluation, I obtained repeat-annotations from reference (1), listed in GenBank accession number AF125183. In agreement with that report, my analyses identified two classes of unique repeats (A and B), positioned with respect to the TSS of the human *H19* gene (Fig. 1). These two repeat classes are within a duplicated DNA segment: repeat unit 1 and repeat unit 2, reference (1). Unit 1 is proximal to the *H19* transcription site and includes A1, B1, B2, and B3; Unit 2 is upstream of the first unit and includes A2, B5, B6, and B7; between the two units, there is a partial repeat (B4), designated B4p in Fig. 1. In initial pairwise alignments, I noted that several of the previous assignments were accurate, *vis-à-vis* start and end positions of the following pair of sequences: A1 and A2; B1 and B5; B2 and B6 (Appendix A). However, even though from results of BLAST, one could deduce that B7 was a partial repeat (1), Clustal O alignments of B3 and B7 partial suggested that B7 corresponded to a somewhat full repeat (see the 2^nd^ and the 3^rd^ pages in Appendix A). Specifically: the start position of B3 in repeat unit 1 shares a stretch of sequence similarity with the corresponding region in repeat unit 2; likewise, the end position of B3 in repeat unit 1 shares a stretch of sequence similarity with the corresponding region in repeat unit 2.

**Fig. 1.**
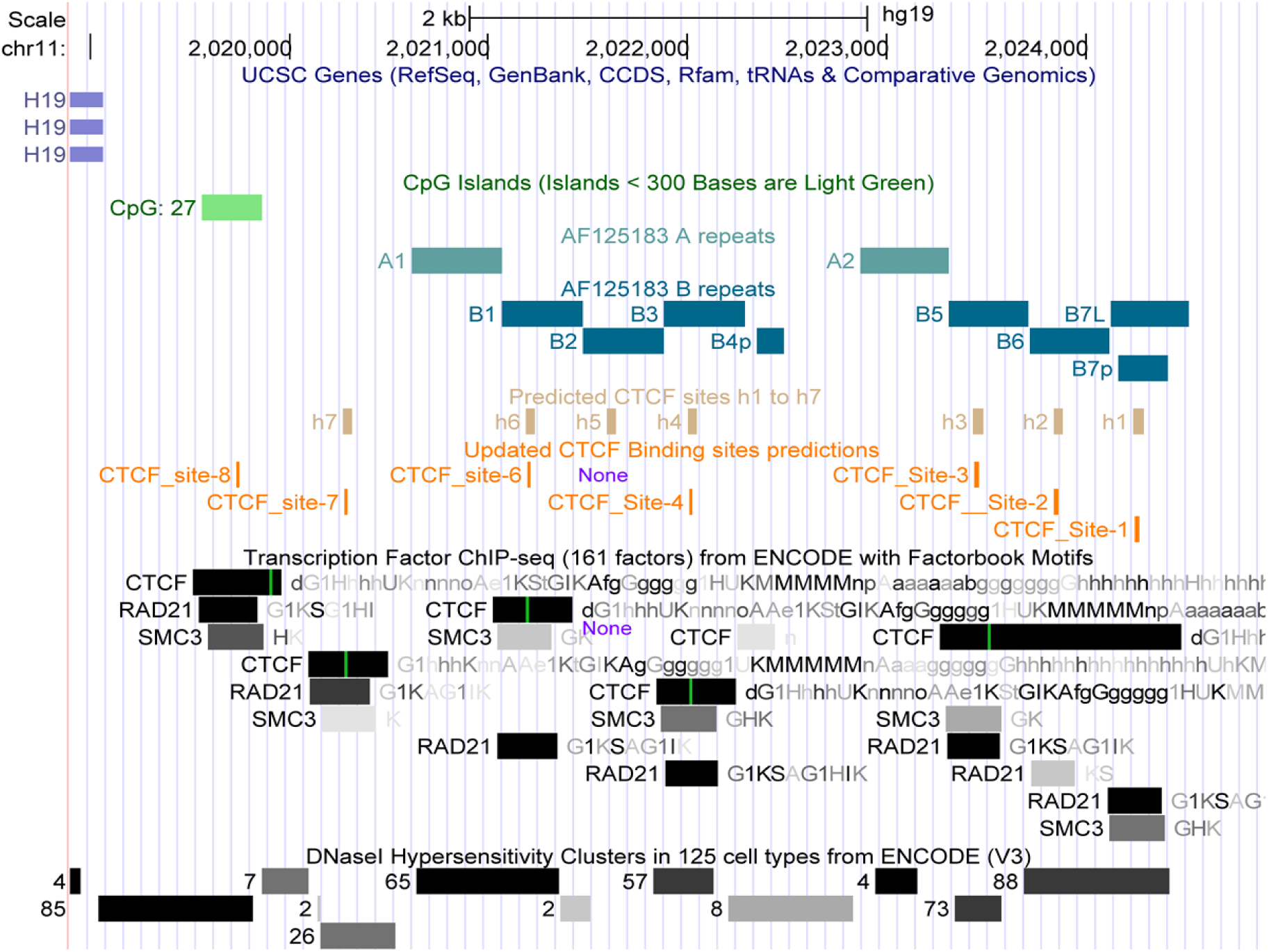
A snapshot from the UCSC genome browser displaying the positions of A and B repeats with respect to *H19* TSS. This figure shows the positions of h1-h7 reported in reference (5), my predicted CTCF sites in the context of results of the ENCODE ChIPs (reported for CTCF, RAD21, and SMC3), and DNase I hypersensitive clusters. Note that results of ChIPs did not find any association of CTCF with h5 (previously reported as CTCF site 5). Also note the position of a previously unknown chromatin boundary (consisting of CTCF, RAD21, and SMC3) in results of the ENCODE ChIP assays reported for the build hg19 of the human genome (17). This boundary includes a predicted CTCF site (designated site 8).

To further evaluate the accuracy of B7 corresponding to a partial repeat, I performed Dotpath analysis. In this and related methods, a program creates a two-dimensional matrix for a pair of sequences, placed along the X and Y axes (Fig. 2). Dotpath simultaneously scans both sequences in a specified window. In the program’s output, sequences that match appear as dots (22). Homologous sequences produce a diagonal line in the center of the matrix; mismatches, including insertions and deletions, cause disruptions in the diagonal. Overall, results of Dotpath analysis of B3 against the long version of B7 (B7L) gave a nearly diagonal line; a few mismatches are detectable in the vicinity of the center of the diagonal (Fig. 2); offset from the diagonal is due to a small deletion within B7L, producing a sequence somewhat shorter than the one obtained for B3 (Appendix A and Table 1).

**Fig. 2.**
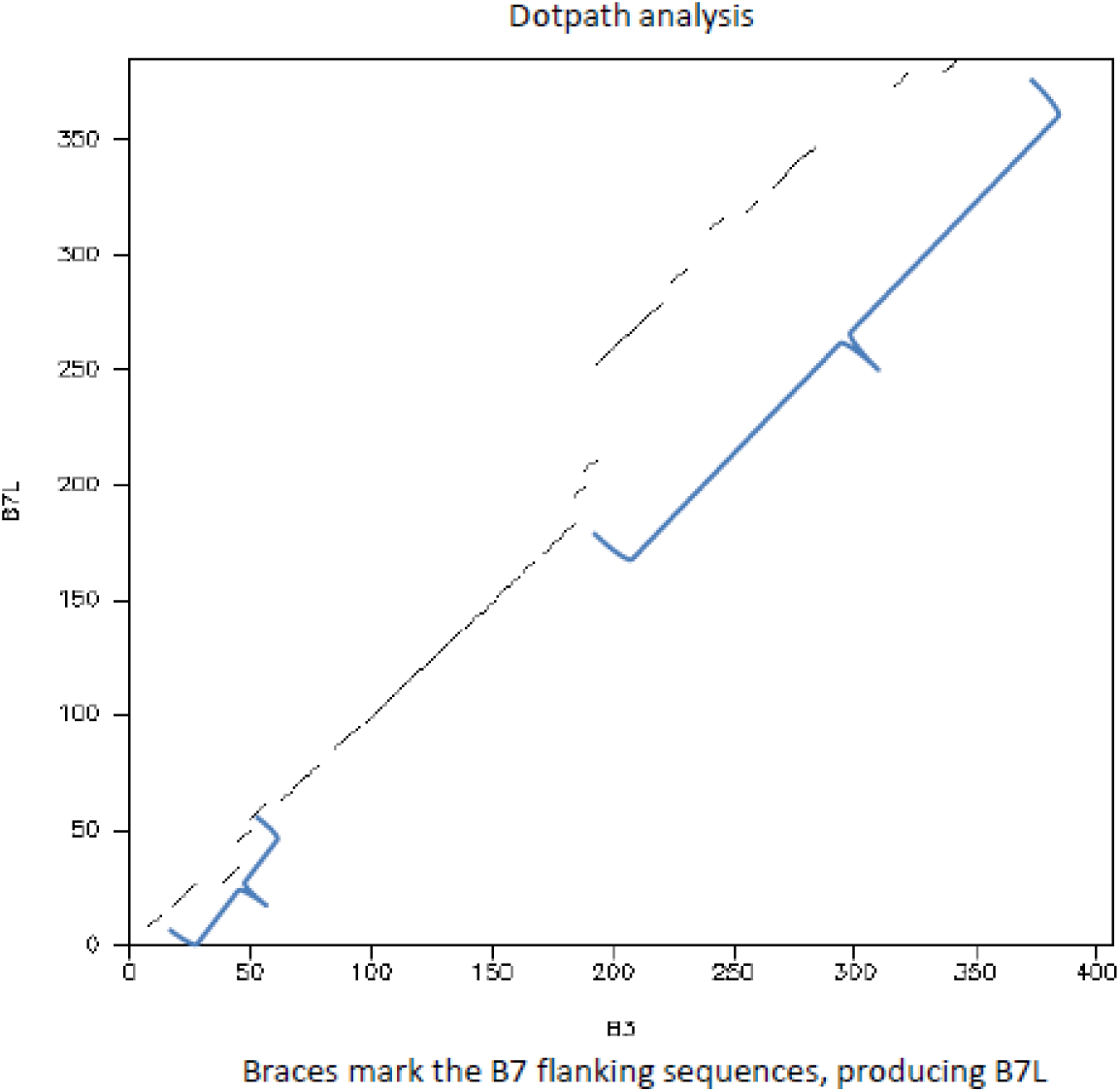
Result of Dotpath analysis indicates that B7 is probably not a partial sequence. Two braces mark the position of sequence similarities obtained from analyses done using Clustal O and Dotpath. These similarities were not detected in results of studies detailed in a previous report (1).

### Multiple sequence-alignment analyses

Comparative sequence-alignments of B repeats could help with examining regulation of the *H19* ICR-mediated transcription and pinpointing the molecular basis of reported genetic abnormities, including Beckwith–Wiedemann syndrome. Towards this goal, I performed several sets of CLUSTAL O alignments of files created to include various combinations of sequences corresponding to B repeats. An initial set contained B1, B2, B3, B4, B5, B6, B7p, and B7L (the longer form of B7). This set was done to determine the missing part in B4 and to display the sequence of B7L with respect to its shorter form obtained in a previous annotation (1). I found that the section missing from B4 is relatively long and corresponds to a 3’ end deletion, producing B4p (Appendix B). Next I examined a set consisting of B1, B2, B3, B5, B6, and B7L while omitting B4p and B7p (Appendix C). This analysis was done to determine the extent of similarity of B7L to B1, B2, B3, B5, and B6. At first glance, it appeared that the 5’ end of B7L shared a limited similarity to the corresponding region in other repeats. However, upon close inspections, I noted several identical nucleotides in this section of B7L to the 5’ end of other repeats (shown in bold in Appendix C). Furthermore, in a set that does include B7L, it is evident that the 5’ end sequences of other repeats vary greatly and thus share a limited similarity (Appendix D). Also, CLUSTAL O alignments located the position of a small internal deletion in B7L, detected in results of Dotpath analyses (Fig. 2 and Appendix C); likewise, CLUSTAL O alignments found a 4 base-pair deletion in B5,as well as 1-base insertions/deletions at various repeat-positions (Appendix C). Thus, overall, results of both Dotpath and CLUSTAL O indicate that B7L is nearly a complete repeat, even though it contains a relatively short internal deletion. This deletion explains why B7L is somewhat shorter than B1, B2, B3, B5, and B6 (Table 1).

### Evaluation of predicted CTCF binding sites

The transcription factor CTCF is an 11 zinc-finger DNA-binding protein that exerts maternal effects on transcription during mammalian oocyte growth, and plays important roles in meiotic maturation and early embryonic development (23). Association of CTCF with a subset of cohesin subunits may contribute to the topological domain architecture and to the formation of regions that exhibit unusually high levels of local chromatin interactions (24). A study predicted 7 CTCF binding regions (h1 to h7), within a segment upstream of the human *H19* gene (10), summarized in (Fig. 3). In the *H19/IGF2* imprinted domain, CTCF binding sites are thought to contribute to formation of chromatin boundaries in the maternal allele (5,9,11). Notably, many publications have relied on the predicted CTCF sites in studies of both Beckwith–Wiedemann and Silver–Russell syndromes. Because of the importance of CTCF to the regulation of the *H19* - *IGF2* imprinted domain, I wished to evaluate the genomic positions of h1 to h7, in the context of the ENCODE data: including the position of DNase I HS clusters (HS-Cs) in chromatin and results of ChIPs obtained for CTCF and two of the cohesin subunits (Rad21 and SMC3). ENCODE (V3) reported DNase I HS-Cs for 125 human cell types (25). The chromatin section that encompasses the *H19* TSS, includes several prominent DNase I HS-Cs: *i.e*. HS-C 85, 65, 57, 4, 73, 88, and 10 (Fig. 1); a numerical value indicates the number of cell-lines containing a HS-C mapping to a specific genomic location. HS-C 85 encompasses the *H19* TSS and extends to include part of a CpG island (CpG27); HS-C 65 overlaps with A1 and B1; HS-C 57 overlaps with B3; HS-C 4 is within A2; HS-C 73 is within B5; HS-C 88 overlaps with B7L and encompasses B6 (Fig. 1). HS-C 10 is further upstream (not shown). Notably, I did not find any HS-C associated with B2, which includes h5, corresponding to the predicted CTCF binding site 5 (Fig. 1).

**Fig. 3.**
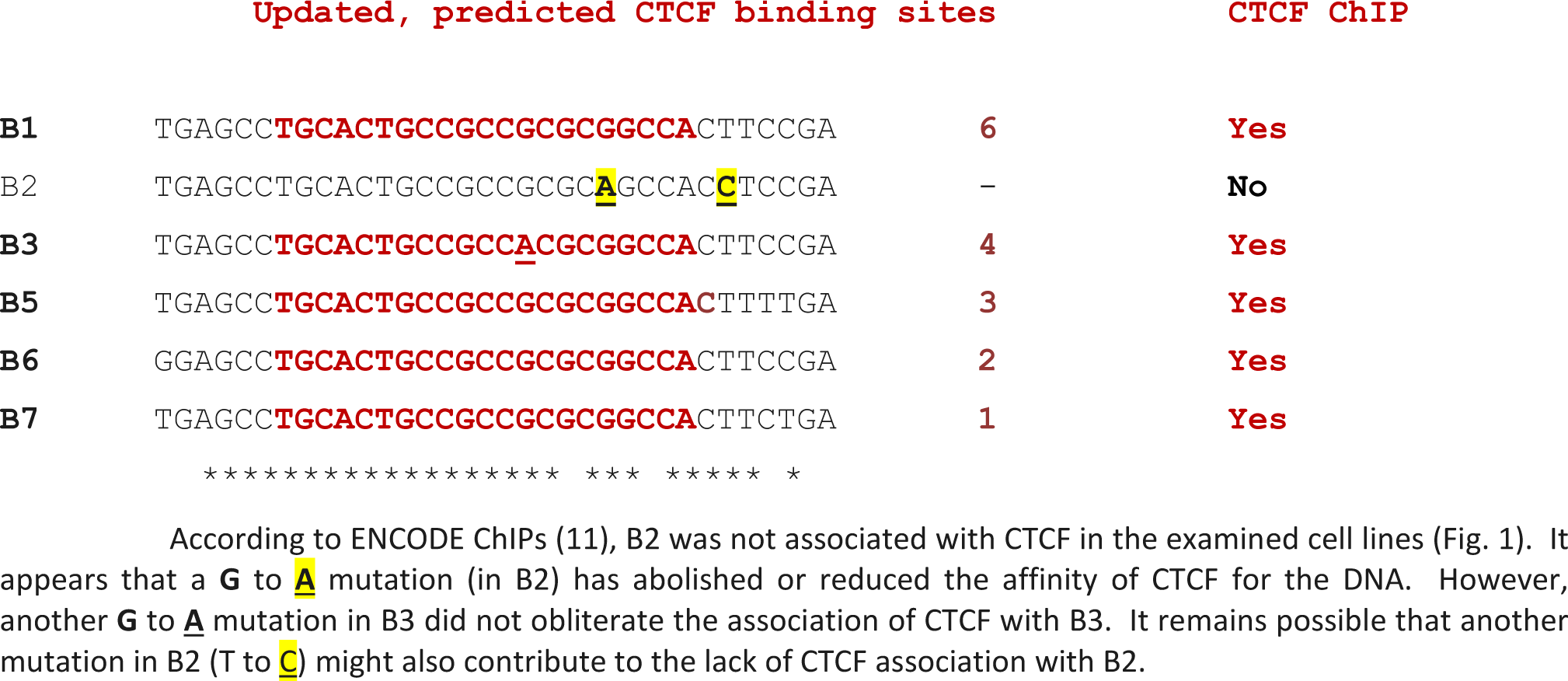
CLUSTAL O alignment of updated, predicted CTCF binding sites in the human *H19* ICR Sequences are shown in the direction found in the chromosomal DNA

The inconsistency noted above led me to closely inspect the genomic positions of h1 to h7 with respect to results of the ENCODE ChIPs, focusing on data reported for CTCF and two cohesin subunits. I also analyzed sequences upstream of the *H19* gene, for potential CTCF binding sites, using a webserver developed and maintained by Yan Cui's Lab, at the University of Tennessee Health Science Center (26): http://insulatordb.uthsc.edu/. Fig. 1 shows CTCF binding sites predicted by that server; they are displayed on a track labeled ‘updated CTCF Binding sites predictions’. With respect to the ENCODE data, CTCF binding sites 1 to 3 map to a relatively broad CTCF-associated region (Fig. 1). ChIP data further revealed that both site 1 and 3 were associated with the cohesin subunit RAD21; site 3 might also be associated with SMC3 (Fig. 1). Sites 4, 6, and 7 are primarily associated with CTCF and RAD21 (Figs. 1 and 3). Thus, among the previously predicted CTCF sites only h5 (in repeat B2) did not map to a CTCF-bound region, according to results of the ENCODE ChIP assays (Fig. 1). To obtain clues into the reason for a lack of CTCF association with h5, I inspected the sequences within the B1 to B7 repeats (Appendix C), summarized in Fig. 3. A close-up view of Clustal O alignment revealed nearly 100% identity in the repeat-sequences encompassing predicted CTCF binding sites. Within that range I noted a G to A mutation in repeat B2, the previously predicted CTCF site 5 in h5; another mutation in B2 (T to C) might also contribute to or responsible for the lack of CTCF association with B2. This interpretation agrees with results of the ENCODE ChIP assays (Figs 1 and 3). Noteworthy could be clinically associated SNPs including rs431825167 and rs483353061 (both in B1) and rs431825169 (in B2).

Notably, results of the ENCODE ChIPs also revealed that a CpG island, upstream of the *H19* gene (CpG27), includes a previously unknown chromatin boundary consisting of CTCF, RAD21, and SMC3 (Fig. 1). Within that region, the Yan Cui's server (26) predicted a CTCF binding site with a DNase I HS cluster: HS-C 85 (Fig. 1). I named the site (CTCF site 8). Clearly, a lack of CTCF association with h5 and the discovery of a chromatin boundary within CpG27, entail mechanistic implications regarding how formation of loops and topological domains would regulate allele-specific gene expression from the *H19 - IGF2* imprinted domain.

## Appendix A

output of CLUSTAL O (http://www.ebi.ac.uk/Tools/msa/clustalo/), analyzing 2 large genomic DNA sections that included the reported A and B repeats. This appendix includes pairwise comparison of the reported repeats units 1 and 2, the reported positions described in the GenBank accession number AF125183, and my comments (this output consists of 3 pages). Note the extend of sequence similarities and the position of insertions and deletions (INDELS)

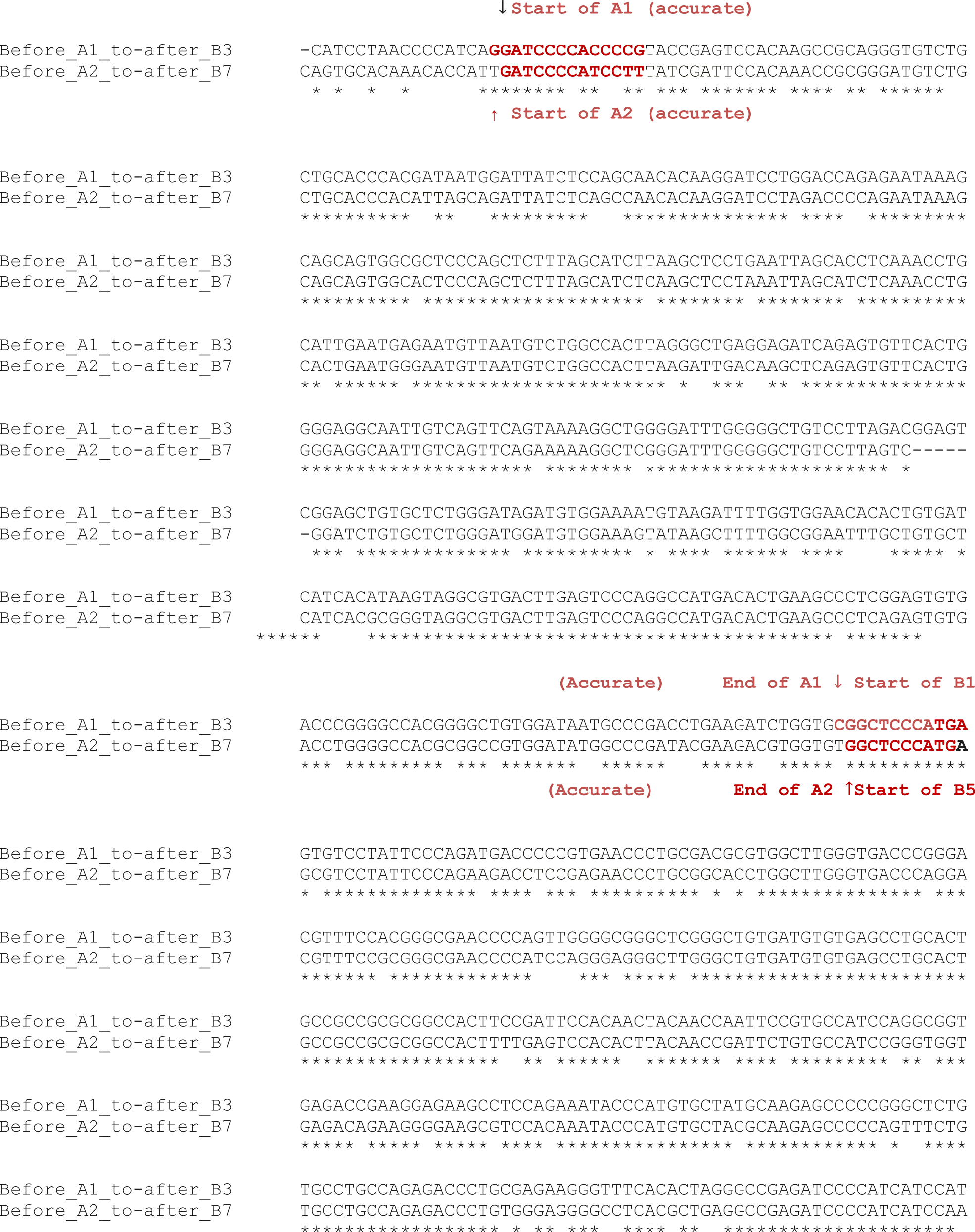
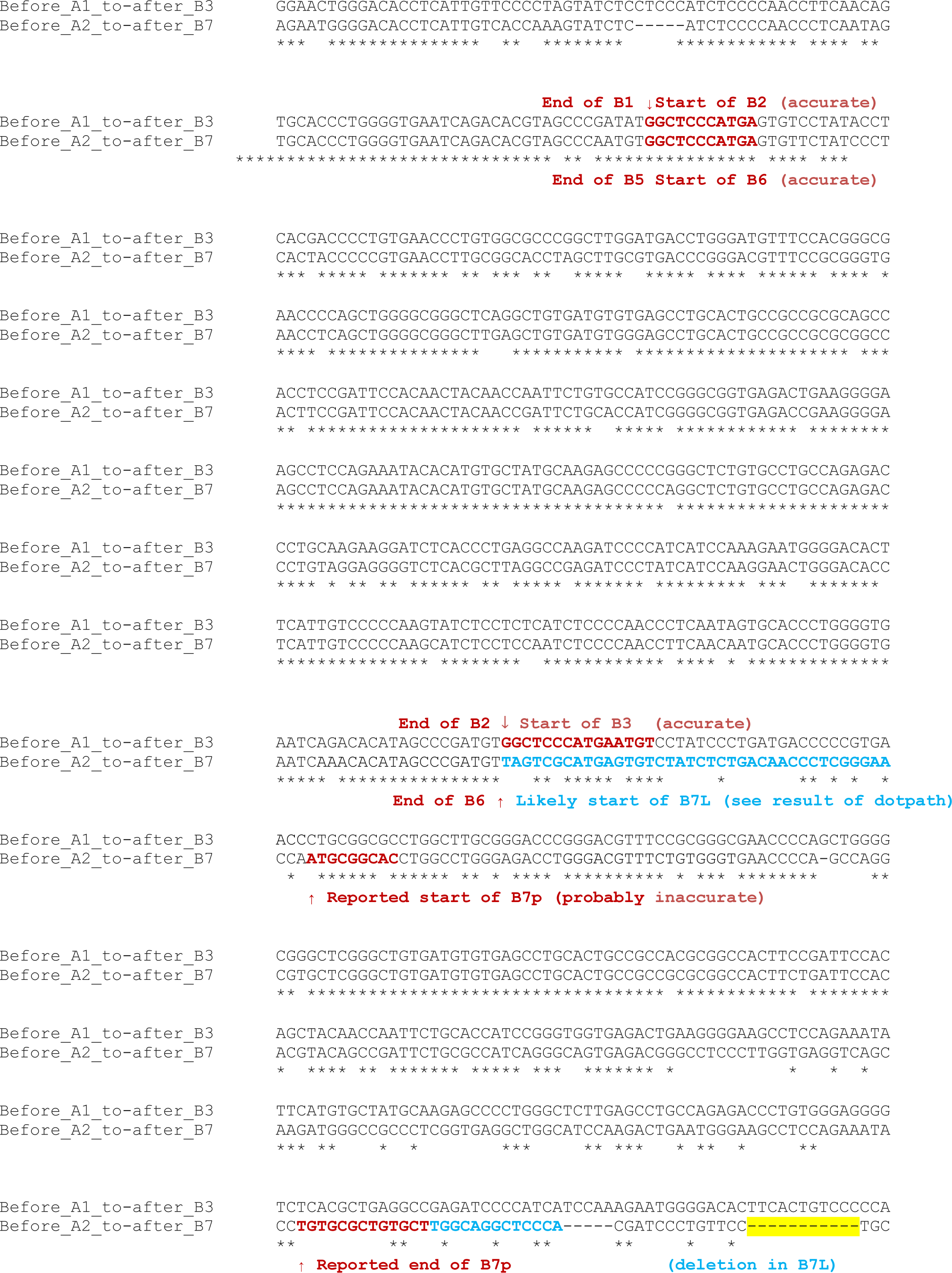
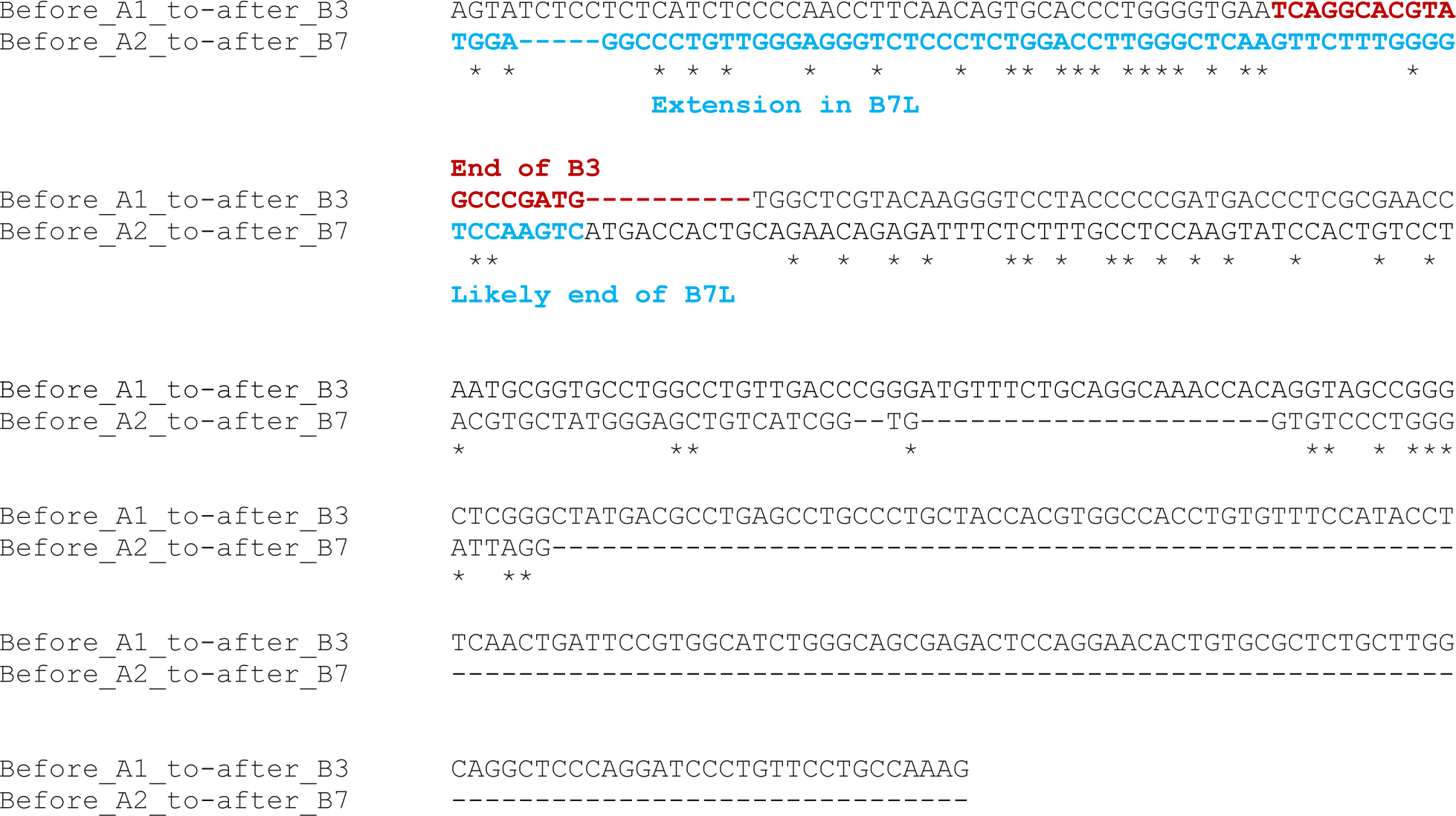

## Appendix B

Alignment of B repeats including B4 partial (B4p), B7 partial (B7p), and the longer form of B7 (B7L). Shown in blue are sequences that are part of B7L but not present in B7p

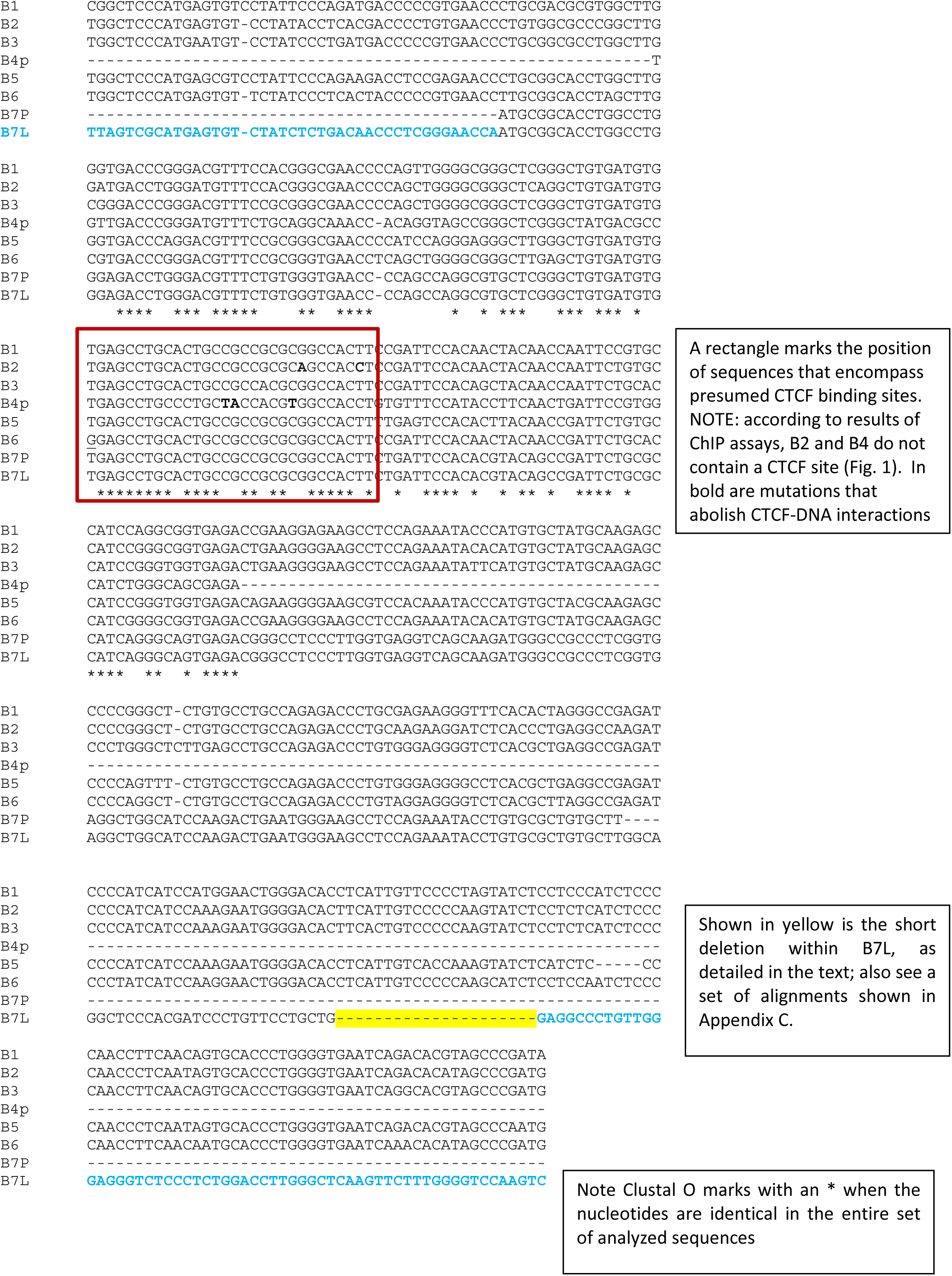

## Appendix C

Clustal O alignments of B1, B2, B3, B5, B6, B7L Highlighted in blue are sequences that were added to B7p, producing B7L

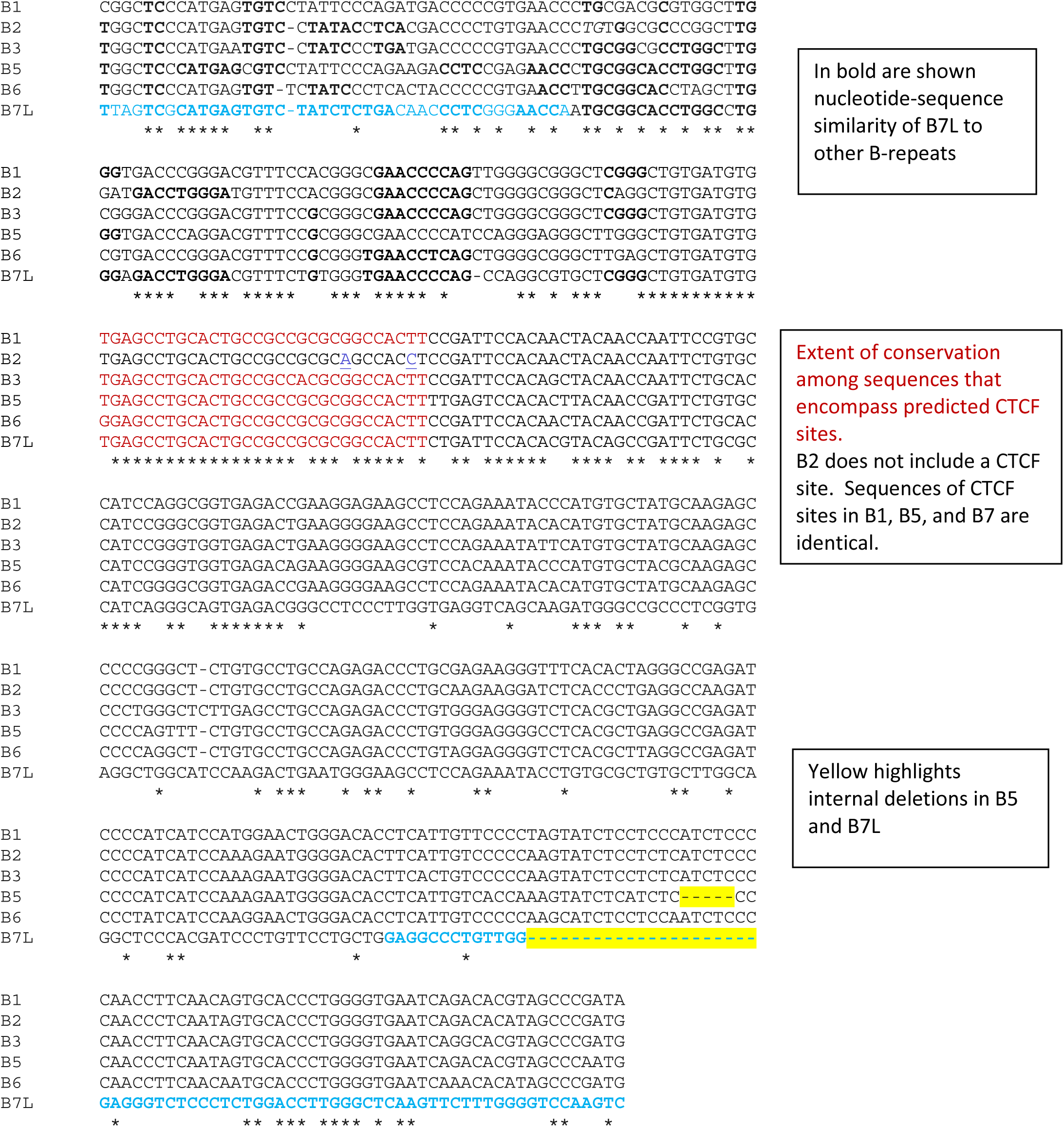

## Appendix D

CLUSTAL O alignment of B1, B2, B3, B5, and B6

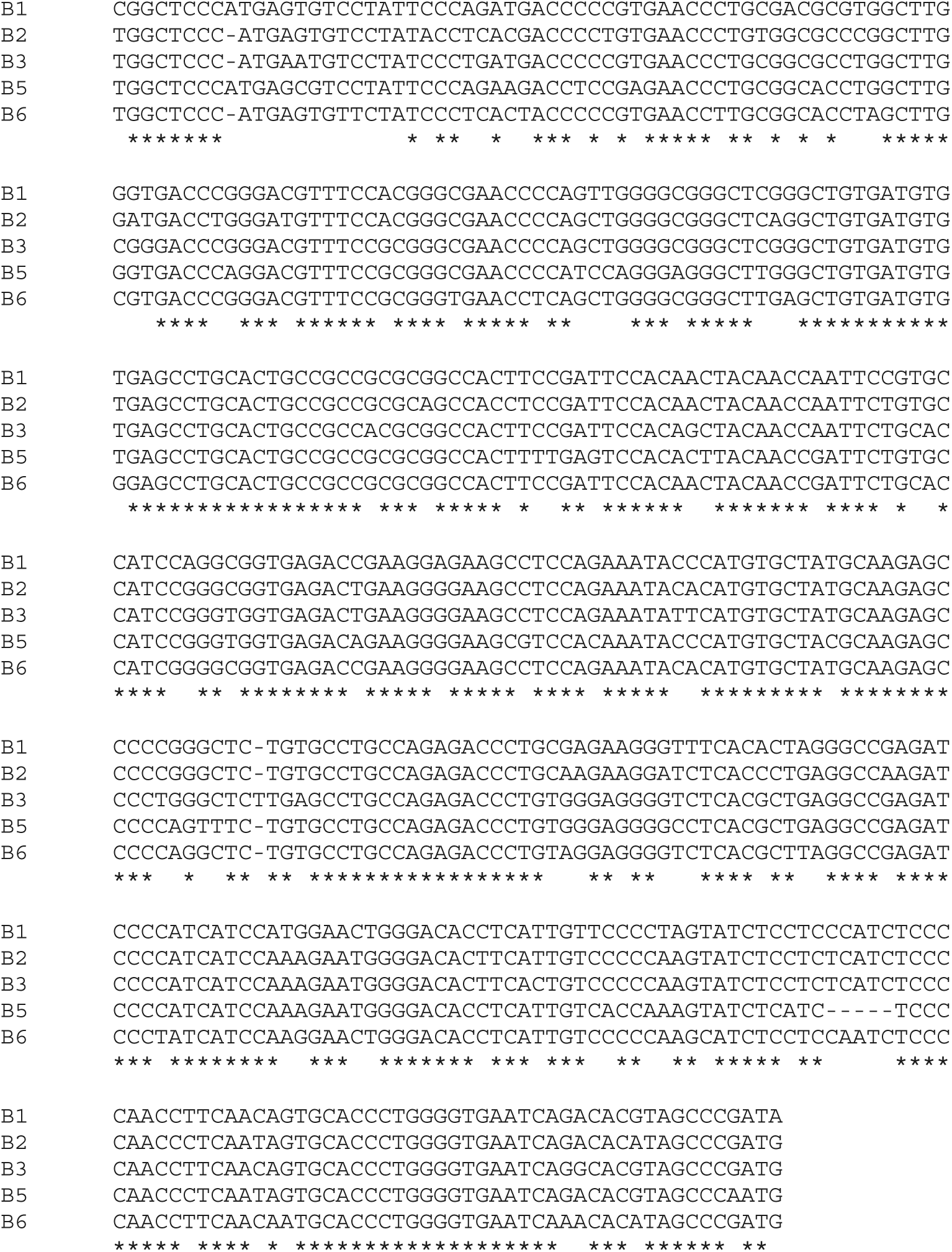

